# Autapses enable temporal pattern recognition in spiking neural networks

**DOI:** 10.1101/2023.11.16.567361

**Authors:** Muhammad Yaqoob, Volker Steuber, Borys Wróbel

## Abstract

Most sensory stimuli are temporal in structure. How action potentials encode the information incoming from sensory stimuli remains one of the central research questions in neuroscience. Although there is evidence that the precise timing of spikes represents information in spiking neuronal networks, information processing in spiking networks is still not fully understood. One feasible way to understand the working mechanism of a spiking network is to associate the structural connectivity of the network with the corresponding functional behaviour. This work demonstrates the structure-function mapping of spiking networks evolved (or handcrafted) for a temporal pattern recognition task. The task is to recognise a specific order of the input signals so that the *Out put* neurone of the network spikes only for the correct placement and remains silent for all others. The minimal networks obtained for this task revealed the twofold importance of autapses in recognition; first, autapses simplify the switching among different network states. Second, autapses enable a network to maintain a network state, a form of memory. To show that the recognition task is accomplished by transitions between network states, we map the network states of a functional spiking neural network (SNN) onto the states of a finite-state transducer (FST, a formal model of computation that generates output symbols, here: spikes or no spikes at specific times, in response to input, here: a series of input signals). Finally, based on our understanding, we define rules for constructing the topology of a network handcrafted for recognising a subsequence of signals (pattern) in a particular order. The analysis of minimal networks recognising patterns of different lengths (two to six) revealed a positive correlation between the pattern length and the number of autaptic connections in the network. Furthermore, in agreement with the behaviour of neurones in the network, we were able to associate specific functional roles of ‘locking,’ ‘switching,’ and ‘accepting’ to neurones.

## Introduction

In biological neural networks, the problem of temporal pattern recognition refers to identifying a sequence of input signals or spikes that carry information. The brain processes such sequences elegantly and responds quickly^1–3^. Biological nervous systems can efficiently differentiate spike patterns distributed across time and space^4^. However, little is known about how individual neurones contribute to processing temporal signals. Interpreting a sequence of spikes generated by a neurone (or a group of neurones) to determine the spatial or temporal structure of a stimulus is a fundamental problem in neuroscience^5^. Analogue sensory information from different modalities, including olfactory, auditory, and visual input, is encoded in the form of spikes^6^, and processed precisely by the brain with incredible speed^7^. The timing of the spikes captures the varying transient intensities of the stimulus, and even a single spike represents remarkable information after stimulus onset^8,9^. These results suggest that temporal features of spikes can precisely represent a stimulus and convey information in biological and artificial spiking neural networks^10^. In the last 15 years, it has been established that sensory information is represented by the precise timing of spikes in the somatosensory, auditory, visual, and olfactory systems^9,11–17^. Furthermore, several studies have shown that information processing in the nervous system is linked to transitions from one spiking behaviour to another^5,18–23^. However, despite extensive research, the association between structural connectivity and the functional behaviour of neural systems remains unclear. Understanding how spiking neural networks process information to gain insight into the computational capabilities of a biological brain is one of the most challenging problems in computational neuroscience^24,25^. A possible way to study the complex computational processing capabilities of the brain is to artificially produce networks with biologically plausible neurones that can perform a computational task.

From a computational perspective, processing temporal spike patterns is a general computational task performed by the brain^26,27^. There are two common ways of learning to recognise temporal patterns: (i) adjusting conduction delays^28–31^, (ii) selecting conduction delays from a spectrum of existing delays^20,32,33^. Spiking neural networks (SNNs) are documented to differentiate temporal patterns by exploiting different time delays and pathways in the network^6^. In neural systems, time delays can be adjusted at the level of synapse, axon, or soma. Adjusting these delays at one or more levels in the network can uncover features of a signal^34^. On the other hand, it is possible that signals produced at different timings arrive together at the readout neurone, generating a maximal response. This phenomenon is used for edge detection in the visual system^35^. Moreover, delays in the network can also be used to identify keywords in continuous speech^36^. This work provides a new way of learning to recognise temporal patterns by evolving (constructing following the rules borrowed from the natural evolution of population and selection) the topology and connection weights of a population of SNNs, without changing or selecting conduction delays.

Artificial spiking neural networks can process a large amount of data using an efficient spiking communication mechanism among neurones such that information is transmitted only when a neurone spikes^9^. Due to their similarities to biological networks, SNNs are widely used to model information processing in animal brains. SNNs have been shown to be computationally more powerful, especially in terms of speed and accuracy, and can solve problems more efficiently than non-spiking neural networks^37,38^. Furthermore, SNNs are considered robust to noise and damage, and their functionality degrades gracefully^39,40^. To understand the learning and information processing mechanism of SNNs, we first need to obtain networks that can perform a specific computational task. Considering the fact that information received from the majority of sensory modalities is temporal in structure, in this work we evolved the structural topology and connection weights of a population of SNNs using a genetic algorithm to recognise temporal patterns. This task can be accomplished by recurrent connections (a form of memory) or delays in the network^29,41^. In general, a recurrent connection can be of any size. However, the shortest possible recurrent connection observed in the nervous system is a self-connection. Self-connections (autaptic connections or autapses) are recurrent synaptic connections between the axon and dendrites or soma of a single neurone (either excitatory or inhibitory). Discovered five decades ago^42^, autapses can be observed in the mammalian brain in the neocortex, hippocampus, and cerebellum^43^. Recent studies suggest possible roles of autapses in the synchronisation of networks^44^, flexible working memory networks^45^, and coherence resonance^46^. However, their functional role remains unknown^47^.

The computational task for SNNs in this work is to recognise a given subsequence of signals in a continuous stream of input signals. A single *Out put* neurone spikes for the correct input pattern while remaining silent for other input patterns. We show the importance of autapses by revealing their possible functional role in state maintenance—a form of memory in the network. Furthermore, SNNs with autaptic connections tend to evolve a simplified switching mechanism to recognise patterns of lengths three and four^48^. Consistent with the evolved networks, we define rules for constructing the topology of a network by hand to recognise patterns up to length six with six interneurones, demonstrating a perfect linear relation between the number of signals in the pattern and the number of interneurones in the network. We show that autapses are crucial for switching the network between states and maintaining a network state. Furthermore, we indicate that the pattern length to be recognised correlates positively with the number of autaptic connections in the network. More specifically, recognising a pattern of n signals requires a network of n interneurones with n-1 autaptic connections. Finally, we demonstrate that, in addition to the other neurones (N1, *N*2, …), a successful recogniser network must have three specialised neurones: a *Lock*, a *Switch*, and an *Accept* neurone. The activity of *Lock* prevents *Out put* from spiking, except when the network receives the second to last correct input signal and allows the *Out put* neurone to spike in response to the correct last input. The *Switch* neurone is responsible for transitions between the network start state and inter-signal network states. The *Accept* neurone forces spike(s) in the *Out put* neurone if the lock is released by the penultimate signal in the pattern to be recognised and switches the network back to the *Start* state by activating the *Switch* neurone.

## Methods

The SNNs in this work consist of adaptive exponential integrate and fire neurones (AdEx) with a set of parameters that result in tonic spiking when a constant input current is injected into the neurone^49^. Each AdEx neurone has four state variables: membrane potential *V*, excitatory conductance *gE*, and inhibitory conductance *gI*, and adaptation *w*, along with 14 parameters (Table 1):

**Table 1.**
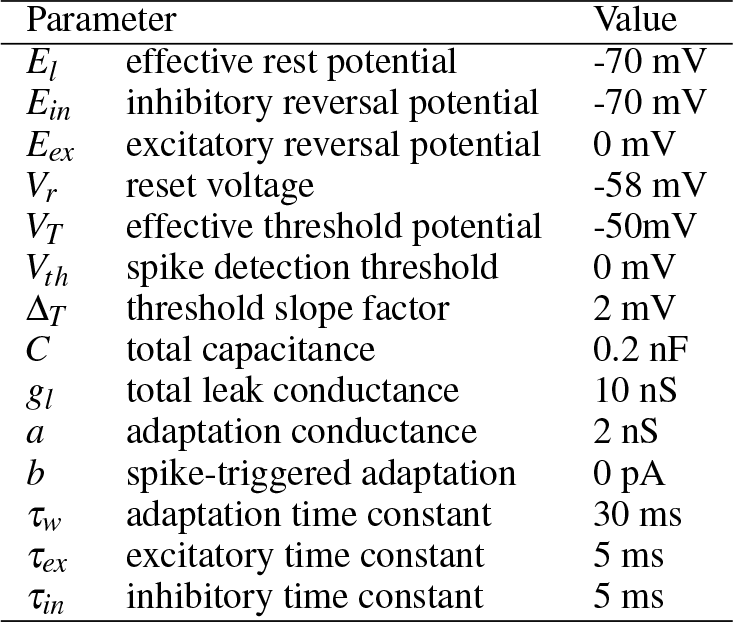
AdEx parameters for tonic spiking.

Out of the 14 parameters, four bifurcation parameters are responsible for the spiking behaviour: the adaptation conductance *a*, the spike-triggered adaptation *b*, the adaptation time constant *τ*_*w*_, and the resting potential *V*_*r*_. The remaining scaling parameters are: the total capacitance *C*, the total leak conductance *g*_*l*_, the effective rest potential *E*_*l*_, the inhibitory *E*_*in*_ and the excitatory *E*_*ex*_ reversal potential, the threshold slope factor Δ_*T*_, the effective threshold potential *V*_*T*_, and the two time constants for excitatory synapses *τ*_*ex*_ and inhibitory synapses *τ*_*in*_.

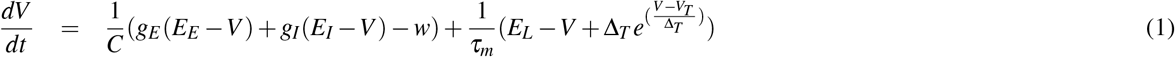

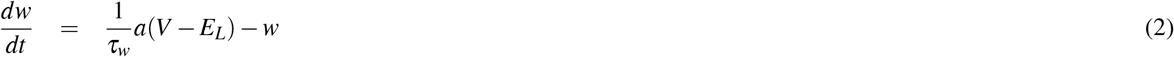

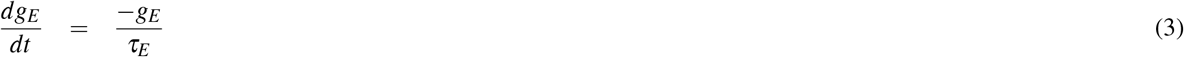

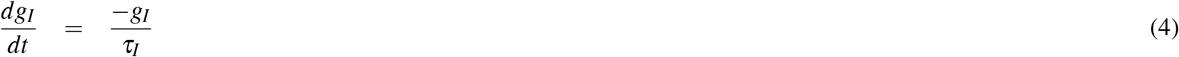

The exponential term in equation 1 defines the spike generation mechanism and the ascent of the action potential. In the mathematical description of the model, a spike is fired at time *t* ^*f*^ when the membrane potential crosses an arbitrary firing threshold value (larger than *V*_*T*_, say +30 mV). When this happens, the integration of the differential equations (Equations 1 to 4) is stopped, the spike time *t* ^*f*^ is recorded, and the voltage is reset to a fixed value *V*_*r*_. This reset describes the descent of the action potential, given by:

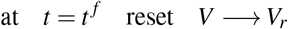

Simultaneously, when a spike is recorded at time *t* ^*f*^, the adaptation current *w* increases by an amount *b*:

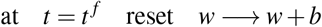

The interaction between the differential equations of the AdEx model and the above two discrete resets can generate a variety of spiking behaviours^49^. In this work, we use the parameters (Table 1) for producing tonic spiking when a step current is injected into a neurone. The state variables are integrated with a time step of 1 ms using Euler integration (using a time step of 0.1 ms or more precise integration algorithms did not affect the results in preliminary experiments, and was not practical from the point of view of computational time, even when using high throughput computing resources made available for this project). The membrane potential is affected by noise; at each time step, a random value is added to it, drawn from a normal distribution with a mean of 0 and a standard deviation of 1 mV.

### Evolution of networks for recognising a pattern of length 3

To obtain networks that recognise a pattern of three signals in a particular order, we used a genetic algorithm originally developed for evolving gene regulatory networks^50,51^, where the topologies of the networks in the population are encoded as linear genomes. Each genome contains a list of genetic elements such that each element has three attributes: a type (input I, output O, dendrite D, or axon terminal A), a sign (+, -), and coordinates (x, y) (Figure 1). A sequence of D elements followed by a sequence of A elements encodes one neurone in the network. To determine the total synaptic strength (weight) of a connection between two neurones in the network, we aggregate the affinity between all A elements of the presynaptic neurone and all D elements of the postsynaptic neurone. Each element has (x,y) coordinates in an abstract affinity space; the smaller the Euclidean distance between two elements i and j, the larger the contribution to the synaptic strength given by 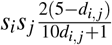 . If the signs (*s*_*i*_ *and s* _*j*_) are the same (different), the contribution is positive (negative). The pre-post connection is established only if the absolute value of the sum of the weight contributions is above a cut-off threshold (to prevent full connectivity); a positive (negative) sum results in an excitatory (inhibitory) connection. In this work, a given presynaptic neurone can excite some of its targets and inhibit others – violating Dale’s principle^52^). However, any violating neurone in the evolved network can be transformed to follow Dale’s principle by dividing it into two parts, one excitatory and one inhibitory (each having the same number of incoming connections, with the same weights, as the original neurone), so that network performance is not compromised^48^.

**Figure 1.**
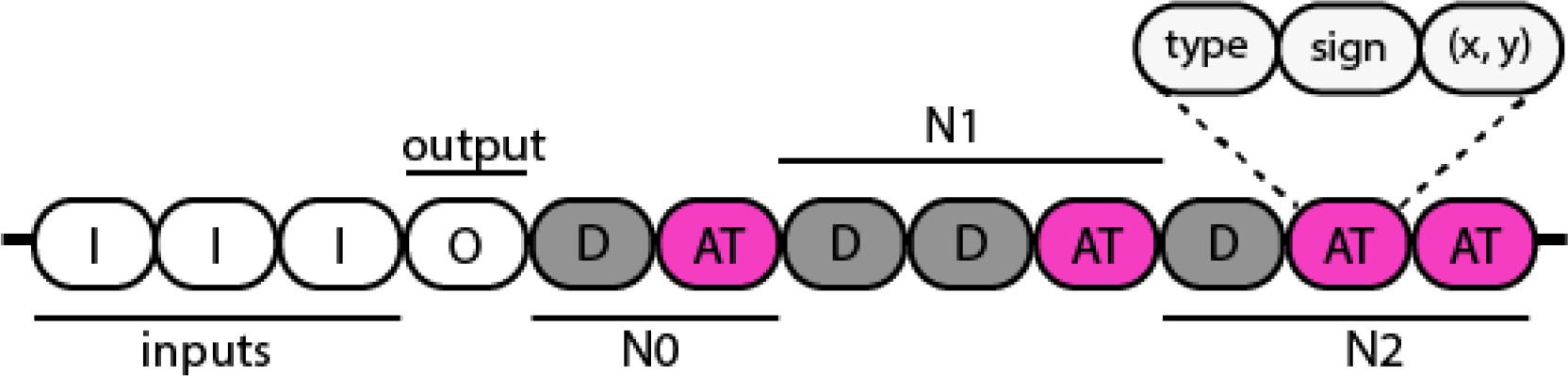
The structure of a random linear genome in the population, encoding a network with three interneurones, three input nodes and one *Out put* neurone. A sequence of D elements (dendrites) followed by a sequence of A elements (axon terminals) is considered one neurone. Each element has a type (I, O, D, A), a sign (+, -) and (x, y) coordinates in 2D space. The strength of a connection between two elements is defined as an inverse function of the Euclidean distance between their coordinates

We use the genetic algorithm^50^ with a constant population size of 300. The initial population is created with random topologies encoded as linear genomes. Subsequent generations are created with a size two tournament selection and an elite count of 10 individuals. Four genetic operators, point mutation, deletion, duplication, and crossover, are described in a previous work^40^, and their probabilities can be found therein. A recurrent SNN for recognising a pattern of three signals was found to require at least three interneurones, one *Out put* neurone and three input nodes. The inputs are not allowed to connect to *Out put* directly, and only the interneurones are allowed to have self-loops (autaptic connections). During evolution, the genetic algorithm can add/remove connections by moving the (x,y) coordinates associated with each genetic element.

The task of the networks is to recognise a pattern of three signals in a continuous stream of signals in which all signals (A, B, and C) occur with equal probability. In an input sequence, the length (duration) of a signal is 6 ms, followed by a silence interval of 24 ms. During evolution, each network in the population is evaluated for six sequences of signals. Four out of these six sequences are generated randomly with an equiprobable occurrence of the three signals (A, B, and C). The remaining two sequences are created by concatenating hard-to-differentiate patterns (ABA, ABB, ABC, BBC) in random order. The fitness function rewards the spiking of the *Out put* neurone in the correct inter-stimulus interval after the occurrence of the correct pattern ABC and penalises spikes in all other intervals. The inter-stimulus interval is defined as the interval between the onsets of two consecutive stimuli.

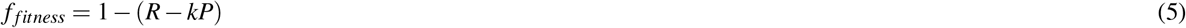

where *R* is the normalised reward given by the number of inter-stimulus intervals after C in which the *Out put* neurone spikes after receiving the correct pattern ABC divided by the total number of correct patterns in the input sequence. The penalty *P* is the number of inter-stimulus intervals in which the *Out put* neurone spikes incorrectly, divided by the total number of inter-stimulus intervals in the sequence (which is one less than the total number of signals in the sequence). As this denominator is large, the normalised penalty *P* is amplified by a constant *k* = 4.

Evolution tends to produce superfluous connections that can be pruned without impairing the performance of the network^48^. To aid the network analysis, we prune the evolved networks by removing a random connection and testing if the performance of the network is compromised. If it is, the connection is put back and labelled vital. Otherwise, the excessive connection is removed from the network. This process is repeated until only vital connections remain in the network.

Pruning revealed structural similarities between the networks obtained from different independent evolutionary runs; thus, the networks that recognised a pattern of three signals were either equal or isomorphic. Moreover, the recognition of a pattern depends on the transitions between the network states. Therefore, the states of an evolved network can be mapped onto the states of the corresponding finite state transducer (FST), accepting a string of three letters.

### Handcrafting networks for recognising patterns of length four and above

Understanding the working mechanism of three-signal networks allowed handcrafting network topologies that recognise longer patterns. For example, a network topology recognising a pattern of three signals can be extended so that it recognises a pattern of four signals by adding a new input, an inter-neurone and six synaptic connections (Figure 2b-c). In the extended network, the input stimuli are renamed *ABCD*. The newly added input A and the neurone *N*4 connect to the existing network; input A connects to *N*4 with an excitatory connection, input *B* (previously named input *A* and connected to neurone *N*3) excites *N*2 (the switch neurone), and input *B* also connects to the newly added neurone *N*4 with an inhibitory connection. The neurone *N*4 connects to *N*3 and to itself with excitatory connections, and the neurone *N*4 and *N*2 (the switch) inhibit each other and thus are mutually exclusive in terms of their activation state. In networks recognising longer patterns, two more connections are essential for seamless switching back to the start state after receiving the last input signal. Therefore, these connections are introduced in networks recognising patterns of length four and above: (i) the last input excites the switch neurone, and (ii) the lock (*N*3) neurone inhibits the *Accept* (*N*1) neurone. These general rules can be used to extend the topology of a four-signal network to obtain the topology for recognising patterns of lengths five and six (Figure 2d-e).

**Figure 2.**
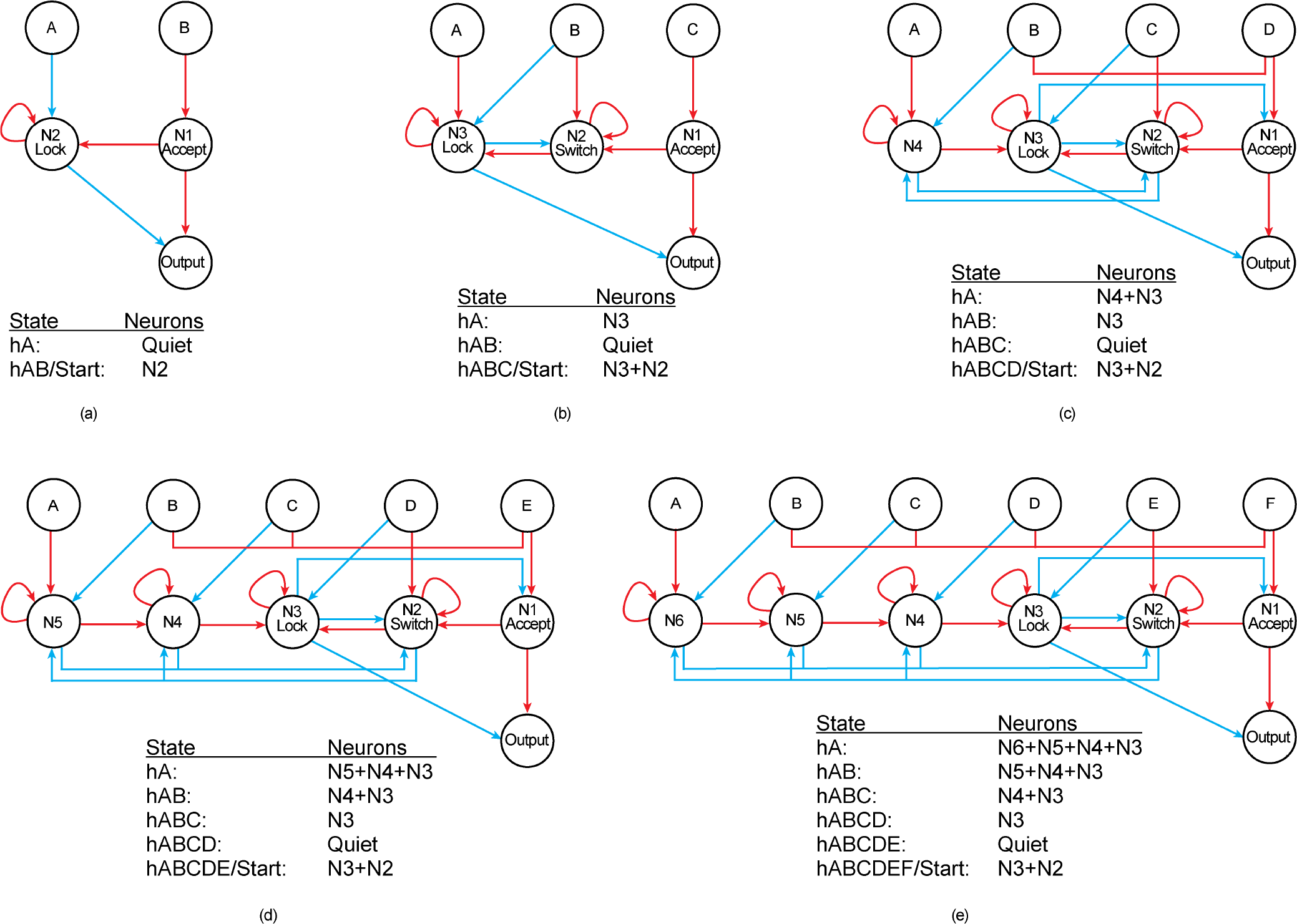
Networks recognising patterns of lengths 2 to 6. The first two networks in panels a and b were obtained with artificial evolution. The table below each network shows transitions between network states, represented by the number of active neurones. The topology is extended by hand to recognise patterns of lengths 4, 5, and 6 (c-e).

The structural topology and the sign of the connections are fixed in the handcrafted networks. We use a genetic algorithm to optimise only the weights of connections. Here, an individual’s genome is the adjacency list of the handcrafted network. In the initial generation, a population of 100 individuals sharing the same structural topology is created. The weights of the excitatory (inhibitory) connections are drawn from a uniform distribution U[0, 10] (U[-10, 0]). The population size is kept constant during evolution with 10 individuals in the elite. The only mutation operator is adding a random number drawn from a normal distribution N[mean=0, SD=1] to the connection weight chosen with a probability of 0.1. Subsequent generations are created with a size two tournament selection and the fitness function is identical to the one for evolving both the topology and the connection weights (Equation 5), rewarding spikes in the correct inter-stimulus interval and penalising spikes elsewhere.

The number of possible patterns increases exponentially with the length of the pattern. There are *n*^*n*^ possible orderings for n signals. For example, a pattern of four signals has 4^4^ = 256 (AAAA to DDDD) possible permutations, and a pattern of six signals has 6^6^ = 46656 (AAAAAA to FFFFFF). This means that when n is large, a given permutation will occur less frequently in a random input stream of signals, and some patterns may not occur at all (so the networks cannot be penalised for recognising them). We have therefore designed a genetic algorithm that runs in two stages. Consider evolving networks for recognising a pattern of length four in a given order. In the first stage, the networks are optimised only for patterns similar to the target pattern ABCD. These similar patterns have the form AXXX, XXXD, where XXX in AXXX (XXXD) is replaced by all 27 possible patterns of BCD (ABC). A sequence of 10,000 signals is created by randomly concatenating the 54 patterns (ABBB to ADDD, AAAD to CCCD). Once the genetic algorithm converges for the first one (finds a network that only responds to the correct pattern ABCD and remains silent for all other patterns), the second stage begins by identifying hard-to-recognise patterns. This requires evaluating the network for a large random sequence with an equiprobable occurrence of signals A, B, C and D. A pattern is considered hard if the network responds incorrectly to multiple occurrences (at least 40% of the total number of occurrences) of a given pattern in a random sequence. Evolution continues with the fitness function in which the penalty coefficient is equal to 50 and the input sequence of 10,000 signals is created by randomly concatenating the 54 patterns (AXXX + XXXD) and hard patterns with equal probability. The second stage (identification of hard patterns, evolution) repeats until the algorithm converges and no hard patterns remain.

## Results

### States of the network correspond to the states of a finite state transducer

To understand how the evolved networks work, we show that the activity of a network can be mapped onto the states of a finite state transducer (FST)^39,48^. An FST is a finite state machine generally used for analysing time-structured data^53^. For example, an FST that accepts a string of three letters needs to have four distinct states (Figure 3b), and so does an SNN evolved to recognise a pattern of three signals (Figure 3a). We established an association between the network states of a pruned network and the states of the corresponding minimal FST. This association can also be obtained for an evolved network, but the mapping is more complex due to superfluous connections^48,54^. The *Start* state of the network corresponds to the continuous spiking of the neurones *N*3 and *N*2 (Figure 3c). Once the network receives the first correct symbol A, it goes into the *hA* (for “had A”) state, maintained by continuous spiking of the *N*3 neurone only. The presence of an excitatory loop on *N*3 preserves the network state until the next signal (A, B, or C) is received. If the network receives another A, it remains in the *hA* state. However, if the network receives a B, it transforms to the *hAB* state retained by no activity in the network (all neurones in the network are quiescent). If C rather than B arrives after A, the network goes back to the *Start* state from the *hA* state.

**Figure 3.**
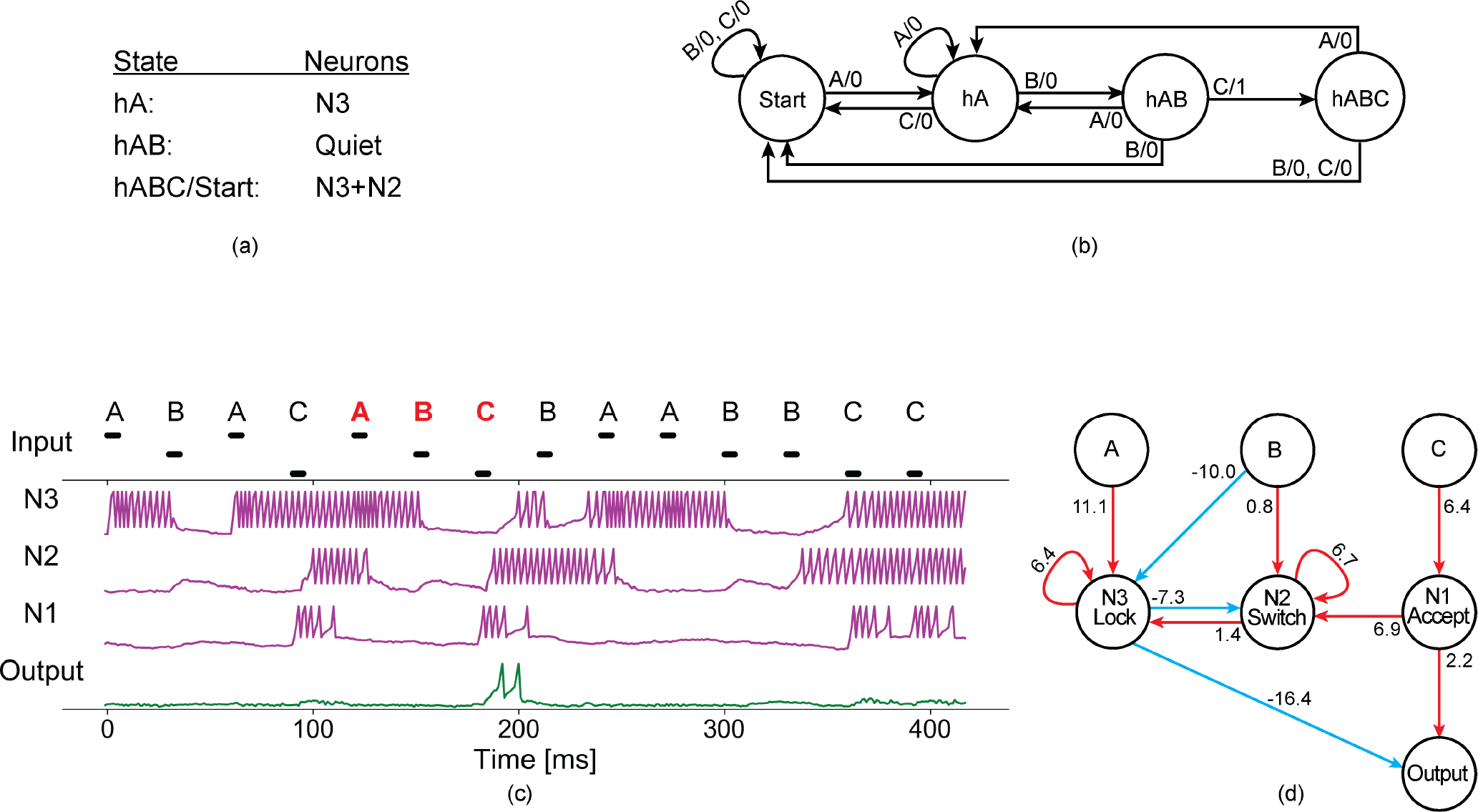
The working mechanism of a network evolved in the presence of membrane potential noise. (a) The states of the network and the corresponding activity of the neurones in the pruned network. (b) The minimal finite-state transducer that recognises ABC. (c) The activity of the network. (d) The topology of the pruned evolved network.

Suppose that the network is in the *hAB* state and a signal C arrives. As a result, the *Accept* neurone spikes several times and activates the *Out put* neurone. The *Accept* neurone also activates the *Switch* neurone to transform the network back into the *Start* state. On the other hand, receiving a signal B when the network is in the *hAB* state activates the neurone *N*2 (explained by the weak excitatory connection from input B to *N*2), which in turn activates *N*3, transforming the network into the *Start* state, represented by continuous spiking of *N*2 and *N*3. Furthermore, if the network receives an A while in the *hAB* state, the network switches back to the *hA* state characterised by continuous spiking of *N*3.

### Specialised role of neurones in the network and the importance of autapses

The pruned networks obtained from independent evolutionary runs for recognising a pattern of three signals were equal or isomorphic^39,48^, allowing us to discover their working mechanism. All resulting networks accomplished the task with four neurones (three interneurones and one *Out put* neurone) and 11 connections (three inhibitory and eight excitatory). Furthermore, networks capable of maintaining network states exhibited self-excitatory loops on at least two interneurones^39,48^. To describe the working mechanism of the networks, first, the role of self-excitatory loops (autapses) is identified. Then, specific roles are associated with the interneurones on the basis of their spiking behaviour. Finally, we demonstrate the contribution of each connection to pattern recognition. The existence of self-excitatory loops enables the evolved networks to maintain network states irrespective of the silent intervals between signals. These recurrent connections permit three active states: the *hA* state by persistent spiking of the *Lock* neurone, and the *hABC/Start* states by causing tonic spiking of the *Lock* and *Switch* neurones (Figure 3a). The only difference between the *Start* and the *hABC* state is the intermittent spiking of the *Out put* neurone. During the evolutionary process, both the *Lock* and the *Switch* neurones form self-excitatory loops with sufficient weights to prevent spiking activity from dying out in the absence of input activity (Figure 3d). In summary, the ability of the network to maintain a state for longer silent intervals requires the formation of autapses in these minimal networks^40,48^.

The interneurones have specialised roles of *locking, switching* and *accepting*. Self-excitation of the *Lock* neurone prevents the *Out put* neurone from spiking, except when the second to last correct input signal (for example, B in ABC) shuts down the *Lock* neurone, enabling the *Out put* neurone to spike for the last correct input signal (C in ABC). If the penultimate correct input signal releases the *Lock*, the *Accept* neurone activates the *Out put* neurone on receiving the last correct input. The *Accept* neurone also sends a signal to the *Switch* neurone, transforming the network back to the *Start* state. The *Switch* neurone is responsible for the transitions between the *Start* state (when *Switch* is active) and the intermediate states *hA* and *hAB* (when *Switch* is quiescent).

### Contribution of each connection to pattern recognition

The contribution of each synaptic connection to the recognition of input patterns is determined by its effect on the behaviour of postsynaptic neurones. The excitatory connection from the input *A* to the neurone *N*3 (*A*→ *N*3) activates *N*3, *N*3 excites itself with an autaptic connection *N*3→ *N*3 and inhibits *N*2 with an inhibitory connection *N*3 → *N*2, thus putting the network in the *hA* state in which only the neurone *N*3 spikes continuously (Figure 3c-d). The continuous spiking of *N*3 also prevents the *Out put* neurone from spiking (due to an inhibitory connection *N*3→ *Out put*). When signal A is followed by B, the inhibitory connection (*B*→ *N*3) forces *N*3 (which maintains the state *hA*) to stop spiking, and the network goes into a quiescent state *hAB*. The input *B* is connected to *N*2 with a weak excitatory connection. This connection prevents an incorrect response of the network to a repeated signal B in the correct pattern ABC. The weight of this connection is adjusted so that the first B cannot activate the *Switch* neurone due to the continued inhibition of *N*3 (*N*3→ *N*2). However, when *N*3 is released by the first B in the correct order, the second B can activate the *Switch* neurone, which in turn activates the neurone *N*3. As a result, the network goes back into the *Start* state (continuous spiking of both neurones *N*3 and *N*2). The positive connection from the input *C* to *N*1 triggers several spikes in the neurone *N*1, and *N*1 passes the activity to the *Out put* neurone. Consequently, the *Out put* spikes in response to the last correct input if the lock has been released by the second to last correct symbol. *N*1 also activates the *N*2*/Switch* neurone, which in turn activates the *N*3*/Lock* neurone, thus transforming the network back into the *Start* state.

### Performance of the handcrafted network for patterns of six signals

The analysis of a network handcrafted for recognising a pattern of six signals shows that the network accomplishes the task of recognising the pattern with seven well-defined network states (Figure 4a). When the network receives the first correct input signal A around 180 ms (Figure 4c), the network goes into the *hA* (had A) state represented by four active neurones, *N*6, *N*5, *N*4, and *N*3. The input *A* is connected directly to *N*6 through a strong excitatory connection, which causes *N*6 to spike on arrival of A (Figure 4b). The self-excitatory loop on *N*6 makes it spike continuously. This spiking activity, in turn, activates *N*5 and prevents *N*2 (the *Switch* neurone) from spiking. The spiking activity of *N*5 persists (due to the self-excitatory loop on *N*5), and *N*5 passes the activity to *N*4, which in turn activates *N*3 (the *Lock* neurone) in a similar way (Figure 4b-c). Meanwhile, when an input signal B is received, the inhibitory connection from the input *B* to *N*6 shuts down *N*6, transforming the network into the state *hAB*, which is characterised by continuous spiking of *N*5, *N*4 and *N*3. The input *B* also excites the *Switch* neurone with a precise connection weight such that only a second B signal can activate the *Switch* neurone and transform the network back to the *Start* state. Similarly, suppose that AB is followed by the third input signal C in the correct order. In that case, the inhibitory connection from the input *C* to *N*5 shuts down *N*5, transforming the network into the state *hABC*, represented by the continuous spiking of neurones *N*4 and *N*3. Next, if the network receives an input signal D in the correct order, it switches off *N*4 through an inhibitory connection from *D* to *N*4, transforming the network into the state *hABCD*, maintained by continuous spiking of *N*3 (the *Lock* neurone) only (Figure 4b-c). These four states from *hA* to *hABCD* are actively maintained by persistent spiking of four, three, two, and one neurone(s), respectively. It is important to note that *N*3 (the *Lock* neurone) is always active, except when the network receives the second to last input signal E in the correct order, allowing the *Out put* neurone to spike for the correct last input signal F. The strong inhibitory connection from *N*3 to the *Out put* neurone prevents *Out put* from spiking for incorrect patterns. If the network is in the state *hABCD* (represented by continuous spiking of the *Lock* neurone) and it receives the second to the last input signal E in the correct order, it switches to a quiescent state by releasing the lock from *Out put* through an inhibitory connection from *E* to the *Lock* neurone (*N*3), which enables the *Out put* neurone to spike in response to the last correct input signal F (Figure 4b-c). The input F activates *N*1 through a positive connection, which in turn passes the activity to both the *Out put* and the *Switch* neurones. The *Out put* spikes in response to receiving the correct pattern ABCDEF, and the *Switch* neurone activates the *Lock* neurone. This transforms the network back into the *Start* state where the *Switch* (*N*2) and the *Lock* neurone (*N*3) spike continuously.

**Figure 4.**
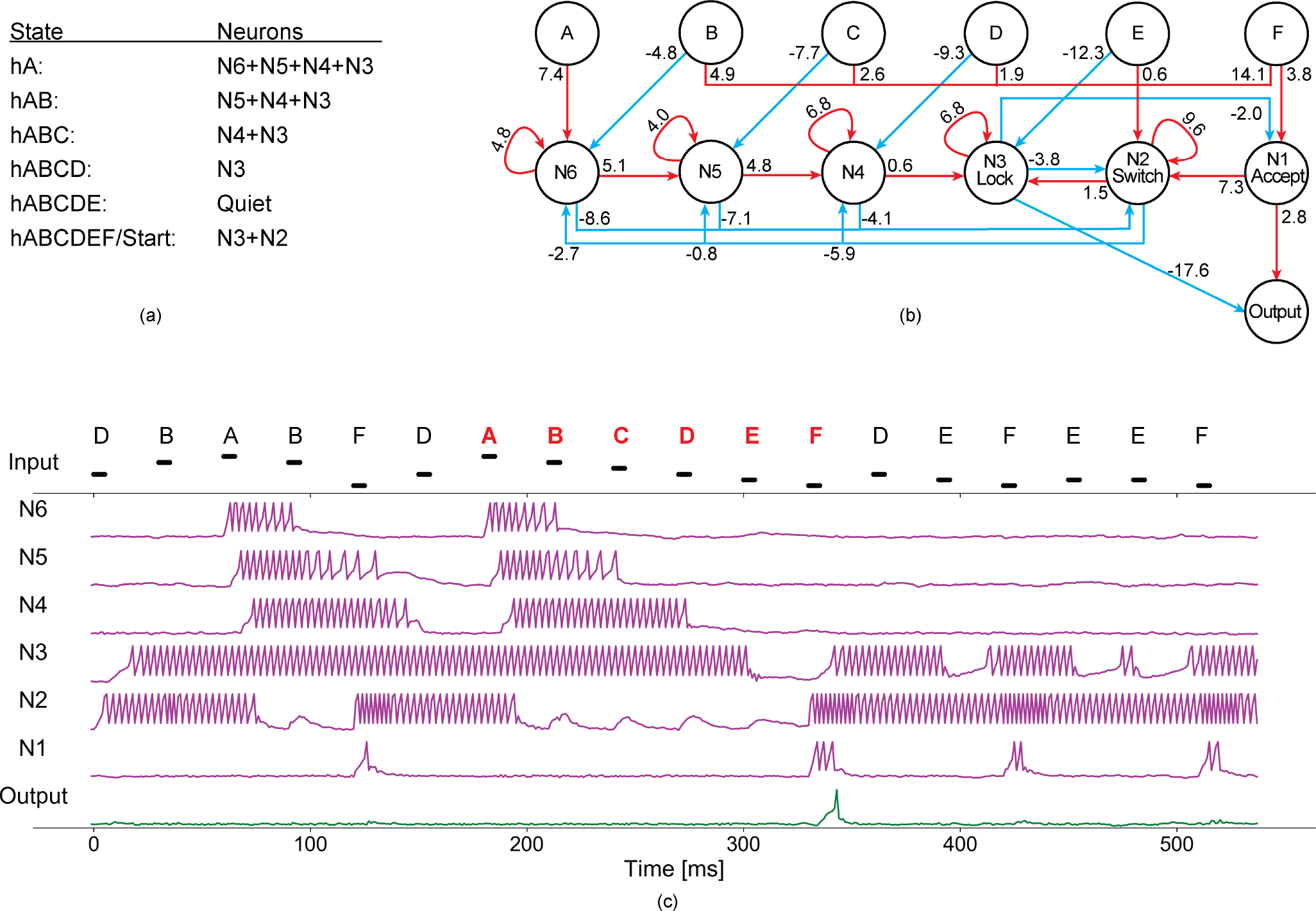
(a) The network states with corresponding active neurones. (b) The handcrafted network for recognising a pattern of length six. (c) The behaviour of each neurone in the network.

The performance of the handcrafted networks is evaluated in terms of precision and sensitivity. Precision is defined as the fraction of *Out put* spikes that are correct, that is, the number of true positives divided by the total number of times the *Out put* spiked, while sensitivity is the fraction of correctly classified target patterns, that is, the number of true positives divided by the actual number of correct patterns in the sequence. The top 10 networks for recognising patterns of lengths three, four, five, and six are re-evaluated for a random sequence of one million signals (Figure 5). The performance of the networks degrades with increasing length of the pattern. Our results show that the precision of the top 10 networks obtained for recognising *ABCDEF* is between 0.73 and 0.96, while for length five (ABCDE) and below the precision is always above 0.94. The networks are then tested with all possible permutations of six signals in six positions with replacement ^6^*P*_6_ (from AAAAAA to FFFFFF), ^6^*P*_7_ (from AAAAAAA to FFFFFFF), and ^6^*P*_8_ (from AAAAAAAA to FFFFFFFF). The testing of the networks with all possible patterns of lengths seven and eight ensures that the performance of the networks is not affected by the proceeding signals, that is, by history. Similarly, all possible permutations of input patterns of lengths three, four, and five are evaluated for up to two preceding signals (Figure 5).

**Figure 5.**
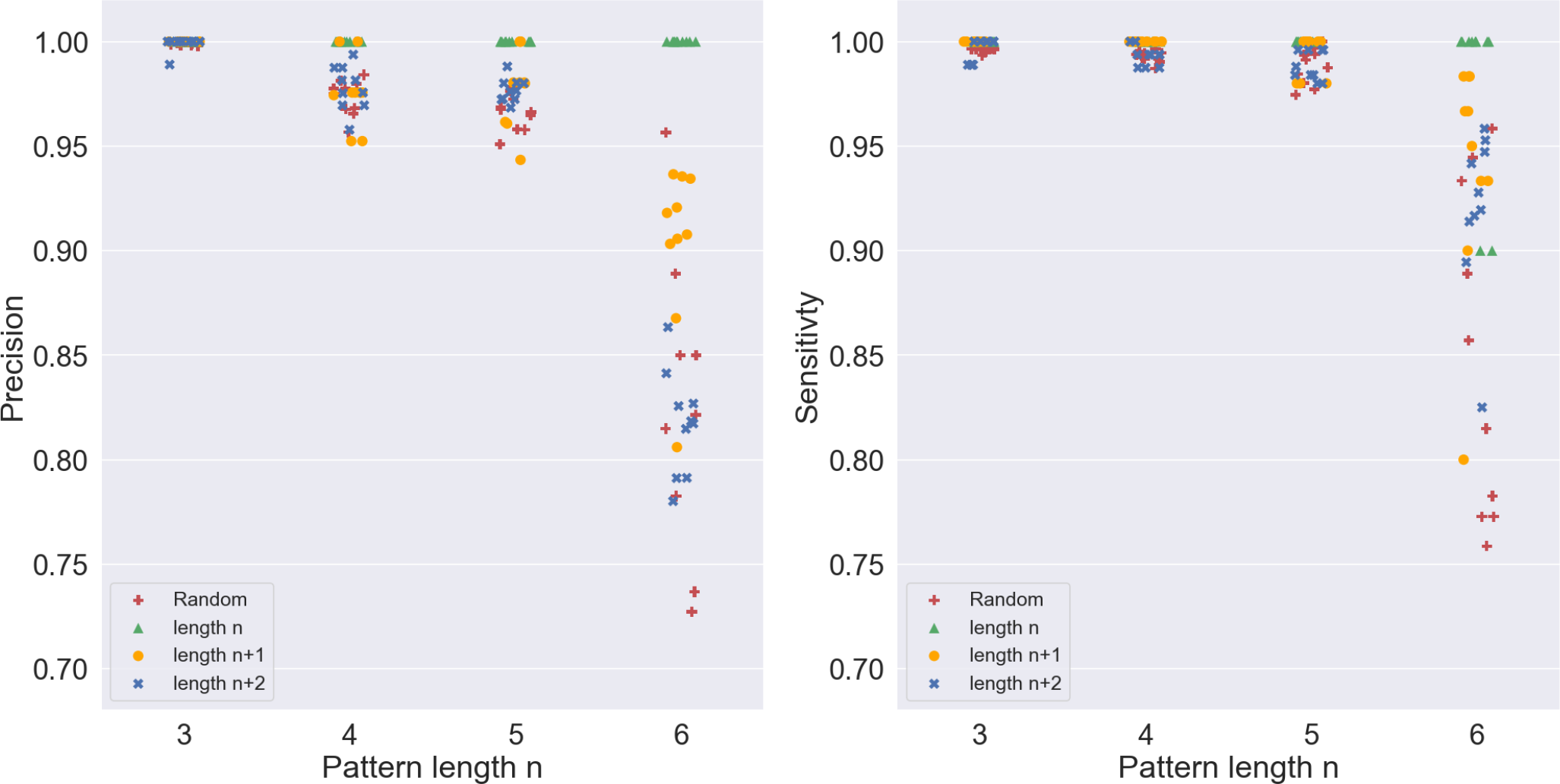
Performance degradation of handcrafted networks with increasing pattern length. The precision (left) and sensitivity (right) of the top 10 networks for each pattern size (3 to 6) are evaluated for a random sequence of length 1 million and all possible patterns of length n, n+1, and n+2.

To demonstrate that all false positives were caused by history or noise, each pattern was presented to the network 10 times. All perfect networks responded to the correct pattern XXABCDEF at least eight out of 10 times, while responding at most four out of 10 times sporadically for a very small number of false positives. This clear margin indicated that handcrafted networks could perfectly recognise correct patterns with up to six signals.

## Discussion

In the present study, we have explored how spiking neural networks can perform temporal pattern recognition tasks without updating or selecting conduction delays. Understanding how the brain processes temporal information is not only an important research topic in computational neuroscience, but also crucial for reproducing artificial networks with brain-like computational capabilities. The neural circuitry of the brain performs a large number of computational tasks, including signal processing, memory, classification, decision-making, etc. It is important to link functional behaviours to the corresponding neural connectivity of the brain. However, the diverse set of computational capabilities, the enormous number of neurones, and the complexity of the brain make a principled mapping of neural connectivity to computational function difficult. In order to make inroads towards a basic understanding of information processing in neural systems, we evolved minimal artificial spiking neural networks for the computational task of temporal pattern recognition.

We found a clear association between structural connectivity and the functional properties of neural networks. We observed that the complexity of the networks increased with the length of the temporal patterns that could be recognised. The number of possible temporal patterns increased exponentially with the length of the patterns, making network training more difficult. Intuitively, the recognition of longer patterns required a larger number of neurones in the network, exponentially increasing the number of connections in the network. Larger and more interconnected networks were harder to train due to the computational cost and larger search space. Consequently, the artificial evolution could not produce an optimal minimal network for recognising a pattern of length four and above, and we resorted to handcrafting the topology of networks for longer patterns (and using artificial evolution only to optimise the weights in these networks).

One of the most salient observations is the emergence of autapses in our spiking neural networks. Although autapses are widely observed in the brain, their functional role in neural information processing is not yet fully clear. Autapses have previously been shown to play a significant role in memory maintenance^55^, oscillatory activity of a single neurone^56^, switching between different network states^57^, synchronisation of network activity^44^, and signal detection^58^. Here, we show the importance of excitatory autaptic connections for network state transitions and network state maintenance. In our work, all active network states are maintained by excitatory autaptic connections, whereas state transitions occur when an excitatory autaptic connection of an active neurone is silenced by a strong inhibitory connection. Furthermore, we examined the relationship between the number of excitatory autapses in the network and the length of temporal patterns that can be recognised. The number of autapses in a network constrains and is equal to the number of distinct actively maintained states in the network. We observed that a minimal network recognising a pattern of length n requires at least n-1 autaptic connections. This is a necessary condition for a network to be able to maintain network states. A sufficient condition is that the weights of the autaptic connections are strong enough to keep the spiking activity from dying out in the presence of longer silent intervals.

Redundant connections in a network have a potential role in learning temporal sequences^59–61^. Although the aim of the present work was to obtain minimal networks for recognising patterns, some redundancy is essential in the network. For example, the extended network that recognises a pattern of length four requires two redundant connections (Figure 2c). If we remove input *A* and neurone *N*4, the four-signal network is reduced to a three-signal network with two redundant connections; an excitatory connection from input *D* to the *Switch* (*N*2) neurone and an inhibitory connection from the *Lock* (*N*3) to the *Accept* (*N*1) neurone. The redundant connection from *D* to *N*2 is required in a four-signal network to activate the switch neurone as soon as the network receives the last correct signal, while the redundant inhibitory connection from *N*3 to *N*1 prevents the *Out put* neurone from spiking when the last symbol D is received in a wrong order. Although the *Out put* neurone is prevented from spiking directly by the *Lock* (*N*3) neurone, this connection reduces the activity that reaches the *Out put* neurone when the lock is active.

Information processing in neural systems is affected by the presence of noise. Noise in the nervous system has previously been documented to play a computational role^62–67^. We previously showed^54^ that networks evolved in the presence of noise are robust to intrinsic (perturbation of parameters) and extrinsic (variation of silent intervals) disturbances. In fact, noise enables networks to maintain network states (a form of memory) when the silent interval between signals is increased during evolution. One of the key findings is that the introduction of noise during evolution simplifies the networks instead of further complicating them. The resulting networks are robust to the removal of connections. Thus, we could prune excessive connections without impairing the network’s performance. As a result, the simplified (evolved and pruned) networks are more efficient and easier to understand. In contrast, the networks that evolved in the absence of noise were fragile and a slight variation of neuronal parameters or network topology can completely alter the functionality of these networks^54^.

The approach of handcrafting the network topology provides insight into neural information processing and the interplay between network connectivity and functional behaviour. However, this work has several limitations that could be addressed in future studies. In particular, the maximum number of active neurones maintaining a network state increases with an increasing length of the patterns that can be recognised. As the number of connections in the network grows, their weights become more difficult to optimise. Due to this limitation, further scaling of the topology produces suboptimal networks. One possible solution to this problem is to build a larger network by connecting two or more three-signal (perfect) networks to recognise more extended patterns. However, the approach of hierarchically connecting smaller networks to build a larger network is not straightforward. We have investigated several ways to interconnect smaller networks to recognise more extended patterns of length 5 and 6, but none of them could outperform the sequential extension of the topologies presented in Figure 2. In future studies, it would be interesting to explore other possible topologies where the maximum number of active neurones (autapses) representing a network state is invariant to the pattern length.

An AdEx neurone can generate a wide range of spiking patterns in response to a step current, depending on the choice of initial parameters^49^. In this study, we are using the parameters for the simplest type of spiking pattern, that is, tonic or regular spiking^49^. A standard leaky integrate-and-fire model (LIF) can also generate this behaviour in response to a step current. In the scope of this study, we do not take advantage of the rich spiking behaviours that an AdEx neurone can have. The neurones in our networks are limited to three possible states: active (tonic spiking), intermittently active, and nonactive (quiet). We observed that an activated neurone may speed up or slow down after receiving an input signal, but we did not take these firing rate changes into account when analysing network behaviour. Identifying network states based on the spiking behaviour of a single neurone would require longer intervals between input signals to notice the difference between their responses. Studying the state transition at the level of a single neurone in the network could result in more efficient solutions. For example, different neuronal behaviours, such as bursting, tonic, and irregular spiking, may represent distinct network states. Moreover, different spiking/bursting frequencies could also represent different network states.

Finally, in this study, we use a genetic algorithm to optimise connection weights in a handcrafted topology. Since all connections in the handcrafted topology are defined according to the state transition table and both the topology and network states are known, a more systematic approach like backpropagation could be employed instead of the genetic algorithm to optimise the weights in the network. This could be done by (i) calculating the errors between the current and the desired network states and (ii) adjusting the connection weights to reach the desired states.

## Conclusions

The present study used a novel combination of evolving and handcrafting spiking neural networks to explore potential mechanisms for temporal pattern recognition in neural systems. Our work demonstrates a link between structural network connectivity and functional network behaviour. A systematic analysis of the resulting networks indicates that excitatory autaptic connections can play a key role in memory maintenance and network state transitions. We predict that the presence of autapses in a neural system implies that the system can perform temporal pattern recognition. A systematic investigation of links between autaptic connections and temporal coding should be an exciting topic for future experimental work.

## Acknowledgements

This work was supported by the Polish National Science Centre (project 2013/08/M/ST6/0092 to B.W.).

## Author contributions statement

All authors contributed equally to the planning of the experiments, the analysis of the results, the editing and the review of the manuscript. M.Y. conducted the experiments using a software platform developed initially by B.W. and other members of his research group.

## Data availability

All data generated or analysed during this study are included in this published article.

